# Direct Small Molecule Modulation of LILRB4 (ILT3) Restores Anti-Tumor Immunity In Vivo and in Patient-Derived Cells

**DOI:** 10.64898/2026.06.10.731269

**Authors:** Somaya A. Abdel-Rahman, Zamara Mariam, Giuseppe Deganutti, Moustafa T. Gabr

**Affiliations:** Department of Radiology, Molecular Imaging Innovations Institute (MI3), Weill Cornell Medicine, New York, NY 10065, USA; Centre for Discoveries in Life Sciences, Coventry University, Coventry, UK

**Keywords:** Immunotherapy, LILRB4, cancer therapeutics, drug discovery, immunomodulators

## Abstract

Small molecule targeting of suppressive myeloid immune checkpoints remains a major challenge in cancer immunotherapy, particularly for non-enzymatic receptors lacking conventional druggable active sites. Leukocyte immunoglobulin-like receptor B4 (LILRB4/ILT3) is an immunosuppressive myeloid checkpoint implicated in tumor immune evasion, T-cell dysfunction, and resistance to immunotherapy across both solid and hematologic malignancies. Here, we report the discovery and characterization of **GL-4512**, a direct small molecule modulator of LILRB4 identified through a Dianthus-based temperature-related intensity change (TRIC) screening platform. Orthogonal biophysical studies, including microscale thermophoresis, surface plasmon resonance, and cellular thermal shift assays, confirmed direct target engagement with nanomolar affinity. Extensive microsecond molecular dynamics simulations combined with site-directed mutagenesis identified a previously unrecognized ligandable pocket within the flexible extracellular domain of LILRB4. Functionally, **GL-4512** disrupted the immunosuppressive LILRB4-SCG2 signaling axis and suppressed downstream SHP1/SHP2 and STAT3 signaling. In patient-derived colorectal cancer and acute myeloid leukemia co-culture systems, pharmacological inhibition of LILRB4 restored anti-tumor immune activity, enhanced IFN-γ and IL-2 production, increased cytotoxic T-cell activation, and reduced tumor-cell viability. **GL-4512** additionally demonstrated favorable pharmacokinetic and safety properties supporting oral in vivo administration. In immunocompetent CT26 syngeneic colorectal tumors, once-daily oral treatment significantly suppressed tumor growth and enhanced intratumoral immune activation. Collectively, these findings establish LILRB4 as a tractable target for direct small molecule immunomodulation and support therapeutic targeting of suppressive myeloid immune checkpoints for cancer using non-biologic modalities.

## INTRODUCTION

Cancer immunotherapy has substantially improved clinical outcomes across a wide range of malignancies, yet many patients either fail to respond or eventually develop therapeutic resistance due to the emergence of highly immunosuppressive tumor microenvironments.^1-4^ Although currently approved checkpoint therapies predominantly focus on T-cell regulatory pathways such as PD-1/PD-L1 and CTLA-4, increasing evidence highlights a critical contribution of suppressive myeloid populations, including tumor-associated macrophages (TAMs), myeloid-derived suppressor cells (MDSCs), and tolerogenic dendritic cells, in restricting effective anti-tumor immunity.^5-9^ These cell populations facilitate immune escape through secretion of inhibitory cytokines, attenuation of antigen presentation, suppression of cytotoxic T-cell function, and establishment of tolerogenic tumor niches.^10-12^ As a result, therapeutic strategies aimed at modulating suppressive myeloid checkpoints have emerged as an important avenue for enhancing responsiveness to immunotherapy and restoring productive anti-tumor immune activity.^13-15^

Among these emerging myeloid immune regulators, leukocyte immunoglobulin-like receptor B4 (LILRB4, also referred to as ILT3) has attracted growing interest because of its prominent role in mediating immune suppression in both solid tumors and hematologic malignancies.^16-20^ LILRB4 is broadly expressed across immunosuppressive myeloid subsets, including TAMs, MDSCs, dendritic cells, and monocytic leukemia cells, where it contributes to maintenance of tolerogenic immune states and inhibition of T-cell activation.^21-24^ Mechanistically, LILRB4 activation initiates signaling through immunoreceptor tyrosine-based inhibitory motifs (ITIMs), resulting in recruitment of the phosphatases SHP1 and SHP2 and downstream suppression of inflammatory immune signaling pathways.^25-27^ Elevated expression of LILRB4 has been linked to poor clinical prognosis, impaired T-cell function, resistance to immune checkpoint blockade, and accelerated tumor progression in multiple cancer settings.^28-30^ Importantly, antibody-based blockade of LILRB4 has shown encouraging preclinical activity, including restoration of T-cell responses, reprogramming of suppressive myeloid populations, and enhancement of anti-tumor immunity.^31-33^ Nevertheless, therapeutic development in this area has remained heavily centered on biologic modalities, whereas direct small molecule modulation of LILRB4 has not been previously explored.

The development of small molecule modulators targeting immune checkpoint receptors represents a fundamental challenge in chemical biology, primarily because these receptors lack conventional enzymatic active sites amenable to activity-based screening approaches. Compared with biologics, small molecules provide several attractive advantages for immune checkpoint modulation, including enhanced tissue and tumor penetration, oral dosing potential, tunable pharmacokinetic behavior, and the ability to access regulatory mechanisms that may not be readily targetable using antibodies.^34-36^ Recent reports, including studies from our group,^37-45^ have demonstrated that protein–protein interactions within the tumor immune microenvironment can be effectively targeted using first-in-class small molecule immunomodulators. Despite this progress, identification of direct LILRB4-targeting small molecules remains particularly challenging because LILRB4 lacks intrinsic enzymatic activity and is therefore not amenable to conventional activity-based screening approaches. To overcome these limitations, we established a Dianthus-based temperature-related intensity change (TRIC) screening workflow designed to detect direct compound engagement with recombinant LILRB4. Candidate hits identified from the primary screening campaign were subsequently advanced through orthogonal biophysical characterization, computational modeling, and functional immune analyses.

In the present study, we describe the discovery of direct small molecule modulators of LILRB4 capable of reversing suppressive myeloid immune signaling and restoring antitumor immune function. Orthogonal biophysical assays validated direct target engagement with high affinity, while downstream functional studies demonstrated modulation of LILRB4-associated signaling pathways and remodeling of suppressive myeloid phenotypes. In patient-derived human immune cell systems, pharmacological targeting of LILRB4 enhanced T-cell activation and reduced tumor-cell viability. Moreover, administration of the lead compound significantly inhibited tumor growth and altered the tumor immune microenvironment in the CT26 syngeneic colon carcinoma model in vivo. Collectively, these findings establish LILRB4 as a tractable target for direct small molecule immunomodulation and support pharmacological targeting of suppressive myeloid checkpoints as a promising therapeutic strategy for cancer immunotherapy.

## RESULTS AND DISCUSSION

### Dianthus Screening Enables Identification of Direct Small Molecule ILT3 Modulators

To identify compounds capable of directly engaging LILRB4, we established a Dianthus-based TRIC screening workflow using recombinant human LILRB4 extracellular domain protein (Figure 1A). Because LILRB4 is a non-enzymatic inhibitory immune receptor lacking conventional catalytic activity, direct target engagement approaches were prioritized over traditional activity-based screening strategies. The Enamine Hit Locator Library, comprising 5,120 structurally diverse compounds, was screened under solution-phase conditions at a final concentration of 10 µM in the presence of fluorescently labeled LILRB4 protein. Compound-associated fluorescence perturbations were quantified as normalized fluorescence changes (ΔF_norm_) relative to vehicle-treated controls.

**Figure 1.**
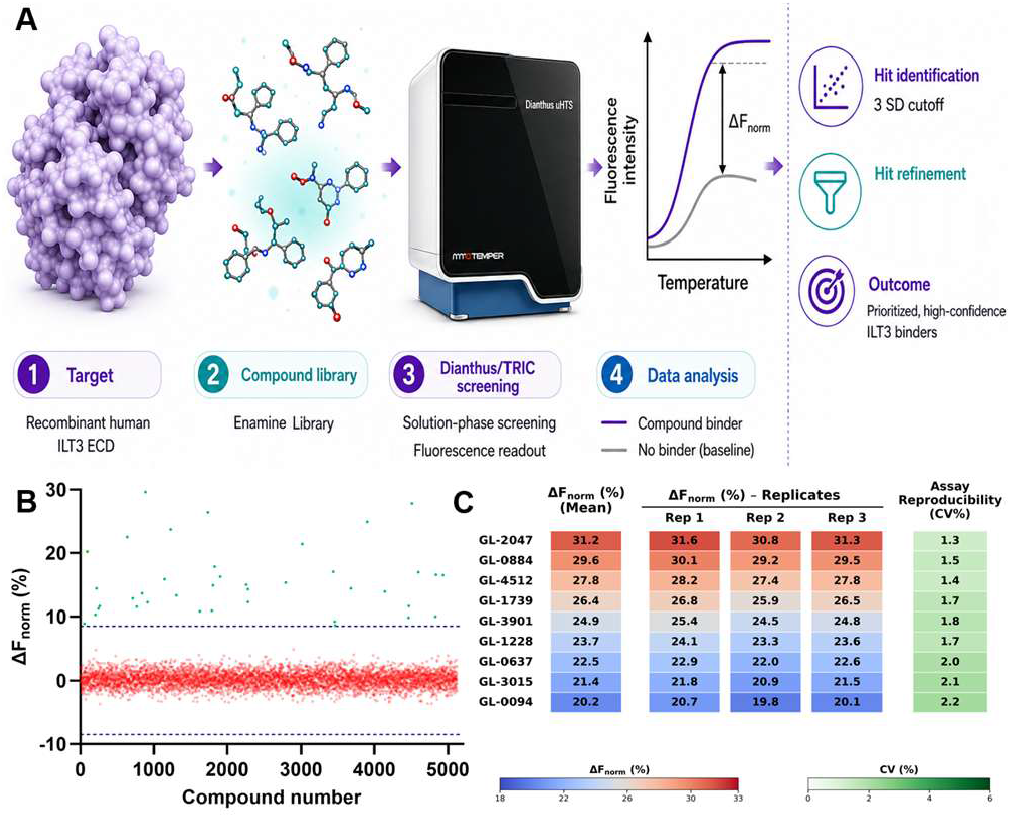
Dianthus/TRIC screening workflow identifies direct small molecule modulators of LILRB4 (ILT3). **(A)** Schematic overview of the Dianthus-based TRIC screening workflow used to identify direct binders of recombinant human LILRB4 ECD. The Enamine Hit Locator Library was screened using fluorescently labeled LILRB4 protein. Compound-induced fluorescence perturbations were quantified as normalized fluorescence changes (ΔF_norm_). Hits exceeding the predefined 3 standard deviation (SD) threshold were advanced for secondary validation. **(B)** Primary Dianthus/TRIC screening results for 5,120 compounds from the Enamine Hit Locator Library. Each point represents an individual compound plotted according to normalized fluorescence response (ΔF_norm_). Dashed lines indicate the ±3 SD cutoff thresholds used for preliminary hit identification. Compounds exceeding the upper threshold were classified as preliminary LILRB4-interacting candidates. **(C)** Secondary validation and reproducibility analysis of prioritized LILRB4 binders. Heatmap representation of the top validated compounds showing mean ΔF_norm_ values, technical replicate consistency, and assay reproducibility expressed as coefficient of variation (CV%).

Primary screening identified 39 preliminary hits that exceeded the predefined hit threshold of 3 standard deviations above vehicle-treated controls, corresponding to an initial hit rate of approximately 0.63% (Figure 1B). Following removal of compounds displaying signal instability, aggregation-associated behavior, or poor reproducibility across replicates, candidate compounds were advanced into secondary validation studies. Heatmap analysis demonstrated strong assay reproducibility and consistent ΔF_norm_ responses across technical replicates for the progressed compounds (Figure 1C). Subsequent validation studies confirmed 9 small molecule binders exhibiting reproducible concentration-dependent interaction profiles consistent with direct LILRB4 engagement, and these compounds were further advanced for orthogonal biophysical characterization.

### Orthogonal Biophysical Studies Validate Direct LILRB4 Engagement by Prioritized Small Molecule Hits

To quantitatively characterize direct binding interactions with LILRB4, prioritized compounds identified from the Dianthus/TRIC screening campaign were advanced into microscale thermophoresis (MST) studies using recombinant human LILRB4 extracellular domain protein. Six hits (Table S1) demonstrated reproducible concentration-dependent binding responses consistent with direct target engagement. Among these compounds, **GL-4512, GL-2047**, and **GL-3901** exhibited the strongest interaction profiles, yielding equilibrium dissociation constants (K_D_) of 18.1 ± 2.4 nM (Figure 2A), 503 ± 19.2 nM (Figure S1), and 3.18 ± 0.65 μM (Figure S2), respectively. The observed MST binding curves displayed clear saturation behavior at higher ligand concentrations together with stable fluorescence baselines across replicates, supporting specific interaction with LILRB4 rather than nonspecific aggregation-associated effects.

**Figure 2.**
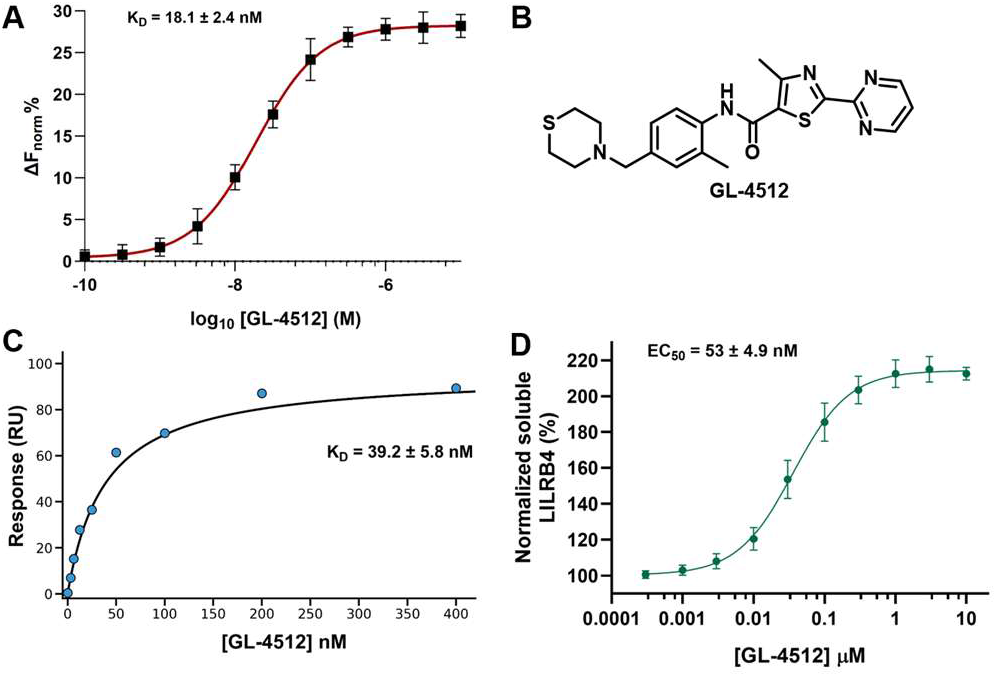
Orthogonal biophysical and cellular target engagement studies validate direct interaction of GL-4512 with LILRB4. **(A)** MST analysis confirming direct binding of **GL-4512** to recombinant human LILRB4 extracellular domain protein. Data represent mean ± SD, n = 5. **(B)** Chemical structure of the lead small molecule LILRB4 modulator **GL-4512** identified through Dianthus/TRIC screening. **(C)** SPR steady-state binding analysis validating direct interaction between **GL-4512** and immobilized LILRB4 protein. **GL4512** exhibited concentration-dependent binding. Data represent mean ± SD, n = 3. **(D)** Dose-dependent CETSA demonstrating cellular engagement of LILRB4 by **GL-4512**. Cells expressing LILRB4 were treated with increasing concentrations of **GL-4512** followed by thermal challenge and quantification of soluble LILRB4 protein. Data represent mean ± SD, n = 5.

Notably, **GL-4512** (Figure 2B) emerged as the highest-affinity binder within the validated series and was therefore prioritized for downstream mechanistic and functional characterization studies. To further confirm direct LILRB4 engagement using an orthogonal label-free biophysical platform, surface plasmon resonance (SPR) experiments were subsequently performed. Consistent with the MST findings, **GL-4512** exhibited concentration-dependent binding to immobilized LILRB4 protein, producing a K_D_ of 39.2 ± 5.8 nM (Figure 2C). The close agreement between MST-derived and SPR-derived affinity measurements provided strong orthogonal validation of direct target engagement. To determine whether the lead compound engages LILRB4 in a cellular context, **GL-4512** was evaluated using a dose-dependent cellular thermal shift assay (CETSA) format adapted from our optimized LILRB4 CETSA platform. Cells expressing LILRB4 were treated with increasing concentrations of **GL-4512** for 60 min, followed by thermal challenge at 51 °C for 3 min and quantification of soluble LILRB4. **GL-4512** induced concentration-dependent stabilization of LILRB4 (EC_50_ = 53 ± 4.9 nM, Figure 2D), supporting direct cellular target engagement and confirming that the compound retains activity in a native cellular environment. Collectively, these findings establish the Dianthus/TRIC workflow as an effective platform for identifying direct small molecule LILRB4 binders.

### Molecular Dynamics Simulations and Site-Directed Mutagenesis Reveal Putative Binding Sites for GL-4512 on LILRB4

To gain mechanistic insight into the molecular basis of **GL-4512** interaction with LILRB4 and to identify potential ligand-binding regions on this highly dynamic immune receptor, we performed extensive molecular dynamics (MD) simulations using the AlphaFold3-predicted structure of human LILRB4. Given the anticipated high flexibility of LILRB4, we favored MD simulations over semiflexible or flexible molecular docking. Two molecules of **GL-4512** (Mol1 and Mol2, Figure 3) were simulated in explicit solvent (i.e., water molecules explicitly represented) under fully flexible conditions, alongside the AlphaFold3 model of LILRB4. This setup should have enabled sampling of putative binding sites for **GL-4512** across four independent replicas of 5 μs each.^46^

**Figure 3.**
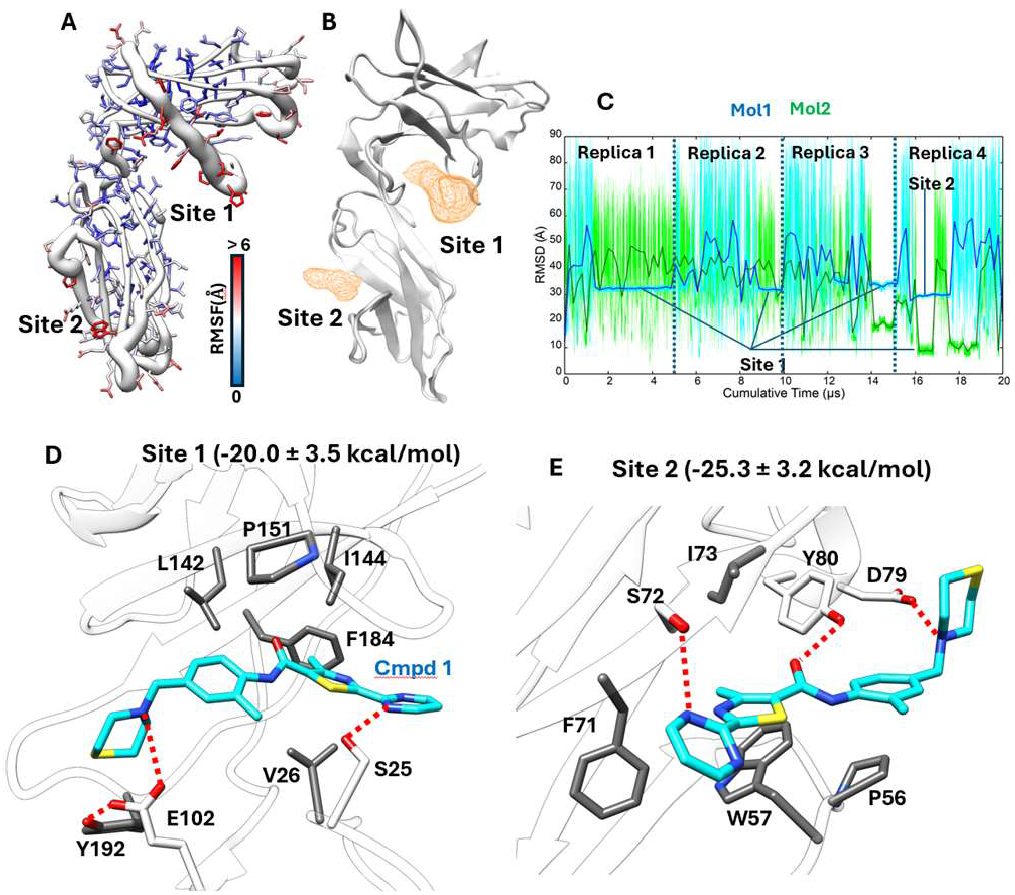
Microsecond MD simulations of GL-4512. **(A)** LILRB4 per-atom RMSF, plotted onto the protein structure. Backbone atoms (white ribbon) diameter is proportional to the RMSF; side chain atoms are colored from white (RMSF = 0) to red (RMSF > 6 Å). **(B) GL-4512** volumetric map (orange volumes, isovalue 0.05) computed for Mol1 and Mol2 over 20 µs of MD simulations, indicating the position of Site 1 and Site 2. **(C)** Mol1 (blue line) and Mol2 (green line) RMSD to the initial coordinates during MD simulations; conformations bound to Site 1 or Site 2 are indicated. **(D)** Representative **GL-4512** binding mode within Site 1; **(E)** Representative **GL-4512** binding mode within Site 2. In **(D)** and **(E)**, hydrogen bonds are indicated as red dashed lines, and residues involved in hydrophobic interactions are in grey stick representation.

LILRB4 exhibited substantial conformational flexibility over the cumulative 20 μs of MD simulations (Figure 3A), as quantified by the root mean square fluctuation (RMSF), which measures per-atom positional variability over time. This behavior is consistent with the experimental requirement to introduce a stabilizing disulfide bond within the D2 domain to enhance structural stability during crystallization (I133C and H143C, PDB 3P2T). Notably, the region corresponding to this flexible D1-D2 hinge, which includes the vicinity of the engineered disulfide in the crystallographic structure, showed the highest engagement with **GL-4512** (designated Site 1; Figure 3B). A second binding pocket (Site 2; Figure 3B) was also sampled and is associated with moderate flexibility, although less pronounced than Site 1 (Figure 3A). Structurally, Site 1 is located between the two immunoglobulin-like domains of LILRB4, corresponding to a flexible hinge region. In contrast, Site 2 is positioned on the opposite face of the D2 domain and is lined by a β-strand.

Across the simulations, three out of four replicas converged with Mol1 bound to Site 1. In Replica 4, Site 2 was occupied for more than 1 μs, indicating a relatively stable interaction. In this same replica, Mol2 also interacted with Site 1. Ligand stability was monitored using the RMSD to the initial configuration (Figure 3C). Within Site 1 (Figure 3C,D), **GL-4512** predominantly formed hydrophobic interactions with residues V26, L142, I144, P151, and Y192, alongside transient electrostatic interactions with S25 and E102. In Site 2 (Figure 3E), **GL-4512** binding was characterized by hydrophobic interactions with P56, W57, F71, and I73, paralleled by electrostatic interactions involving S72, D79, and Y80. Molecular Mechanics Generalized Born Surface Area (MM/GBSA) binding free energy calculations suggest that Site 2 may be energetically more favorable for **GL-4512** binding, with ΔG ≈ −25.3 kcal/mol compared to −20.0 kcal/mol for Site 1. The greater hydrophobicity of Site 1 suggests that favorable desolvation (i.e., displacement of energetically unstable water molecules from the pocket) may contribute significantly to ligand binding.

Given the high flexibility of LILRB4, it is plausible that additional conformational rearrangements are required to fully stabilize **GL-4512** in the bound state and that these rearrangements were not completely sampled within the accessible MD timescale. This limitation is inherent to simulations based on AF3 models rather than experimentally determined dynamic ensembles. However, taken together, our computational results suggest two possible binding sites on the LILRB4 surface. Site 1 appears to be kinetically more accessible but less specific, whereas Site 2 provides a more favorable balance of hydrophobic and polar interactions, consistent with higher druggability.^47^

To experimentally evaluate the putative ligand-binding regions identified through MD simulations, representative residues from each predicted binding site were selected for site-directed mutagenesis followed by MST binding analysis. Since Site 1 is highly hydrophobic, we reasoned that mutation of V26, L142, I144, and P151 to alanine would likely be compensated by neighbouring residues. Thus, we generated the Y192A mutant because of Y192’s dual effect during MD simulations: direct interactions with **GL-4512** and the coordination of E102, which facilitated transient electrostatic interactions with GL-4512’s positively charged thiomorpholine nitrogen atoms (Figure 3D). In parallel, the W57A mutant was generated to interrogate Site 2, which exhibited more favorable MM/GBSA binding energies together with a structurally defined interaction network involving aromatic and polar residues.

MST analysis revealed that alanine mutation of Y192 produced only minimal changes in **GL-4512** binding affinity relative to wild-type LILRB4 (Figure S3), suggesting that although Site 1 is frequently sampled during MD simulations, this region may primarily function as a kinetically accessible encounter surface rather than the dominant productive binding pocket. In contrast, alanine mutation of W57 resulted in a substantial reduction in binding affinity, with an approximately 32-fold increase in the apparent equilibrium dissociation constant compared with wild-type protein (Figure S3). These findings strongly support Site 2 as the principal binding region responsible for stable high-affinity interaction of **GL-4512** with LILRB4. Collectively, the combined MD and mutagenesis studies support a model in which **GL-4512** initially samples multiple transient conformational states on the flexible LILRB4 extracellular surface before stabilizing within a more ligandable and structurally defined pocket centered around W57 within the D2-associated region. The strong sensitivity of **GL-4512** binding to W57A mutation further supports the functional importance of aromatic and hydrophobic interactions within Site 2 and provides experimental validation for the computationally predicted binding model.

### GL-4512 Disrupts the LILRB4-SCG2 Axis and Suppresses Downstream Immunosuppressive Signaling

To determine whether direct engagement of LILRB4 by **GL4512** functionally interferes with ligand-mediated receptor activation, we next evaluated the ability of **GL-4512** to disrupt the interaction between LILRB4 and secretogranin II (SCG2), a ligand implicated in suppressive myeloid signaling and tumor immune evasion. Using a time-resolved fluorescence resonance energy transfer (TR-FRET) competition assay, **GL-4512** produced concentration-dependent inhibition of the LILRB4-SCG2 interaction, with an IC_50_ value of 37 ± 2.8 nM (Figure 4A), supporting direct pharmacological disruption of this immunosuppressive ligand-receptor axis.

**Figure 4.**
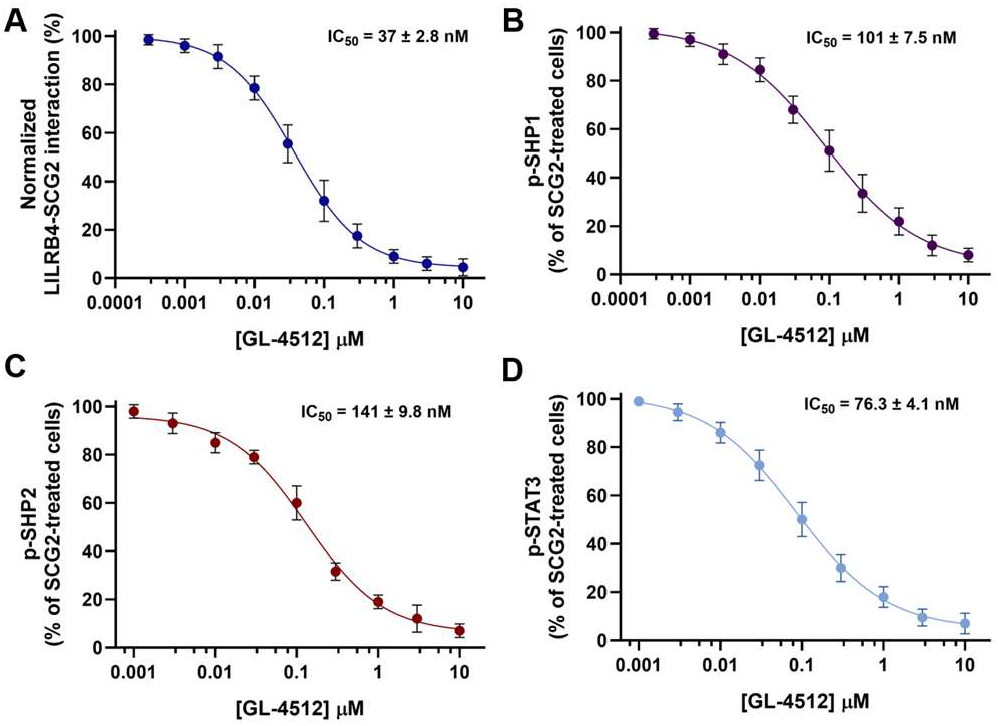
GL-4512 disrupts the LILRB4-SCG2 interaction and suppresses downstream inhibitory signaling pathways. **(A)** TR-FRET competition assay demonstrating concentration-dependent inhibition of the LILRB4-SCG2 interaction by **GL-4512. (B-D)** ELISA-based quantification of SCG2-induced intracellular signaling in LILRB4-expressing THP-1-derived macrophage-like cells treated with increasing concentrations of **GL-4512. GL-4512** produced dose-dependent suppression of **(B)** phospho-SHP1, **(C)** phospho-SHP2, and **(D)** phospho-STAT3 signaling relative to SCG2stimulated cells. Collectively, these findings demonstrate that **GL-4512** functionally disrupts ligand-dependent LILRB4 signaling and suppresses downstream immunosuppressive signaling pathways associated with myeloid immune regulation. Data represent mean ± SD (n=5). Nonlinear regression analysis was used to calculate IC_50_ values.

Because LILRB4 signaling is mediated through recruitment of SH2 domain-containing phosphatases downstream of intracellular immunoreceptor tyrosine-based inhibitory motifs (ITIMs), we next investigated whether **GL-4512** modulates proximal inhibitory signaling events induced by SCG2 stimulation. Exposure of LILRB4-expressing THP-1-derived macrophage-like cells to SCG2 resulted in robust induction of SHP1 and SHP2 phosphorylation, consistent with activation of canonical suppressive LILRB4 signaling pathways. Treatment with **GL-4512** attenuated SCG2-induced phosphorylation of both SHP1 and SHP2 with IC_50_ values of 101 ± 7.5 nM and 141 ± 9.8 nM (Figure 4B,C), indicating effective blockade of proximal inhibitory signaling downstream of LILRB4 engagement.

We next evaluated downstream effects on STAT3 signaling, a key transcriptional regulator associated with suppressive myeloid phenotypes, tumor-associated immune dysfunction, and resistance to anti-tumor immunity. SCG2 stimulation induced substantial phosphorylation of STAT3 in LILRB4-expressing cells, whereas treatment with GL-4512 markedly suppressed p-STAT3 levels in a dose-dependent manner (IC_50_ = 76.3 ± 4.1 nM, Figure 4D). Collectively, these findings demonstrate that pharmacological targeting of LILRB4 by **GL-4512** disrupts ligand-dependent inhibitory signaling at multiple mechanistic levels, including receptor–ligand engagement, proximal SHP phosphatase activation, and downstream STAT3 signaling. These results establish **GL-4512** as a functional small molecule modulator capable of suppressing LILRB4-mediated immunosuppressive signaling pathways relevant to tumor immune evasion.

### GL-4512 reverses SCG2-driven immunosuppression in patient-derived solid and hematologic tumor co-culture systems

To determine whether pharmacological inhibition of LILRB4 by **GL-4512** restores anti-tumor immune function in clinically relevant cellular systems, we next evaluated the activity of **GL-4512** in ex vivo immune co-culture models representing both solid and hematologic malignancies (Figure 5A). In the colorectal cancer setting, primary PBMCs isolated from colorectal cancer patients were co-cultured with HCT116 or HT-29 tumor cells in the presence of SCG2 to induce suppressive LILRB4 signaling. SCG2 stimulation markedly suppressed anti-tumor immune activity, as evidenced by reduced IFN-γ and IL-2 secretion together with increased tumor cell viability (Figure 5B-D for HCT116 cells and Figure S4A-C for HT-29). Treatment with **GL-4512** restored cytokine production in a concentration-dependent manner (0.1 – 10 μM) and significantly reduced tumor cell viability, indicating recovery of immune-mediated tumor killing. At higher concentrations, the activity of **GL-4512** approached that observed with a blocking anti-LILRB4 antibody used as a target-specific positive control (Figure 5B-D for HCT116 cells and Figure S4A-C for HT-29).

**Figure 5.**
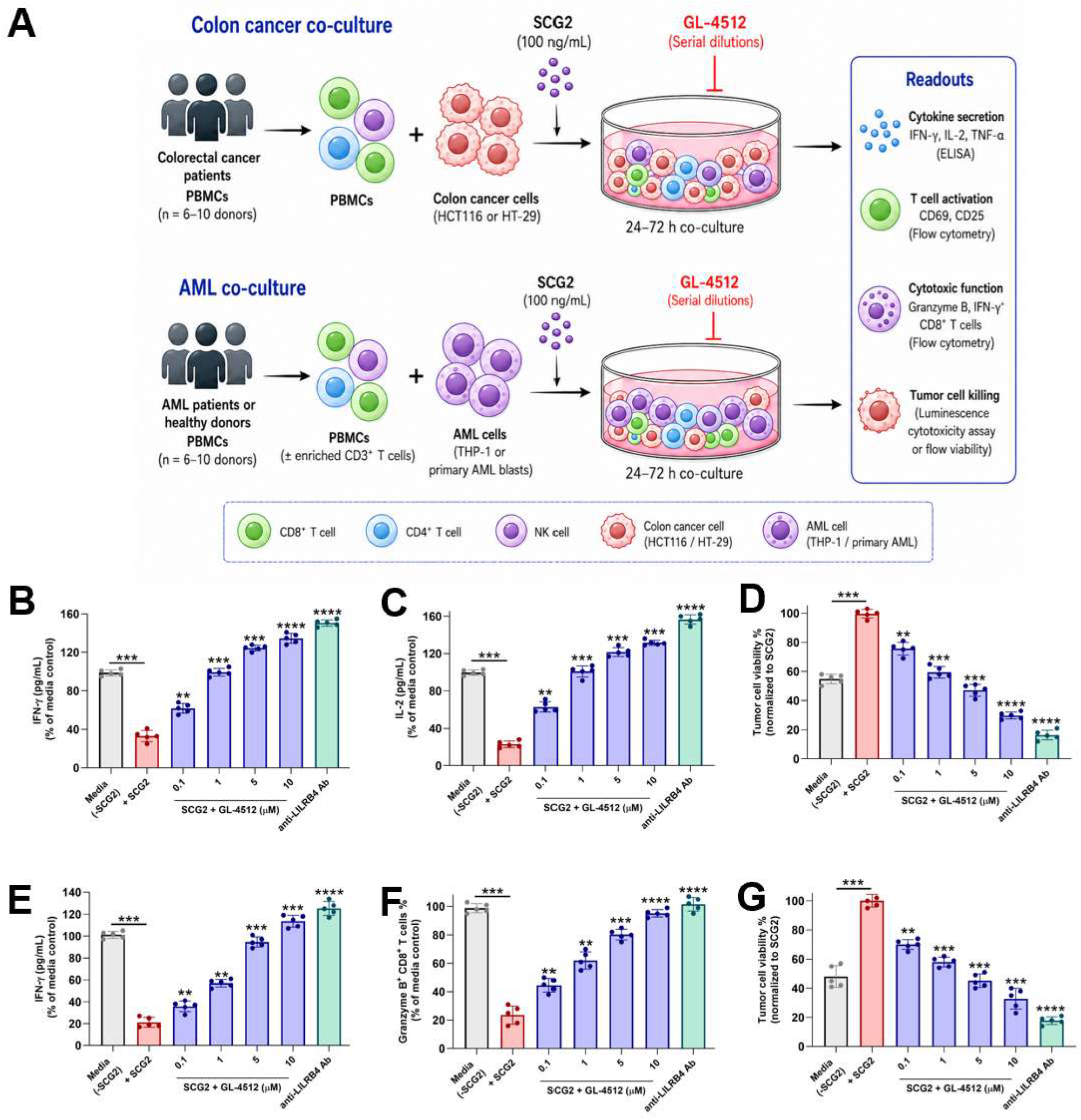
GL-4512 restores anti-tumor immune activity across colorectal cancer and AML co-culture systems. **(A)** Schematic representation of ex vivo co-culture workflows used to evaluate the immunomodulatory activity of **GL-4512** in solid and hematologic tumor contexts. For colorectal cancer studies, PBMCs isolated from colorectal cancer patients were co-cultured with HCT116 or HT-29 cells in the presence of SCG2 and increasing concentrations of **GL-4512** (0.1 – 10 μM). For AML studies, PBMCs from AML patients or healthy donors were co-cultured with THP-1 cells or primary AML blasts under analogous conditions. Functional readouts included cytokine secretion, T-cell activation, cytotoxic function, and tumor-cell viability. **(B**,**C)** In colorectal HCT116 cancer co-cultures, SCG2 stimulation suppressed anti-tumor immune activity, resulting in reduced IFN-γ and IL-2 secretion. Treatment with **GL-4512** restored cytokine production in a concentration-dependent manner. **(D) GL-4512** significantly reduced colorectal HCT116 cancer cell viability in SCG2-treated co-cultures, consistent with restoration of immune-mediated tumor killing. **(E)** In AML primary blasts co-cultures, **GL4512** restored IFN-γ secretion suppressed by SCG2-mediated LILRB4 signaling. **(F) GL-4512** increased the frequency of Granzyme B^+^ CD8^+^ T cells in a dose-dependent manner, indicating recovery of cytotoxic T-cell function. **(G) GL-4512** significantly reduced AML primary blasts viability in SCG2-treated co-cultures. In multiple functional assays, the activity of **GL-4512** approached that observed with a blocking anti-LILRB4 antibody used as a target-specific positive control. Data represent mean ± SD from independent donors performed in technical replicates. Statistical significance was determined using one-way ANOVA with Tukey’s multiple comparisons test. \*\**p* < 0.05, \*\*\**p* < 0.001, \*\*\*\**p* < 0.0001 relative to media +SCG2.

To further assess whether the immunomodulatory activity of **GL-4512** extends beyond solid tumors, we next evaluated the compound in acute myeloid leukemia (AML) coculture systems using THP-1 cells or primary AML blasts cocultured with PBMCs or enriched CD3^+^ T cells (Figure 5A). Similar to the colorectal cancer model, SCG2 stimulation promoted a suppressive immune phenotype characterized by reduced IFN-γ secretion, impaired cytotoxic T-cell activation, and increased AML cell viability (Figure 5E-G for AML blasts and Figure S4D-F for THP-1 cells). **GL-4512** significantly reversed these suppressive effects in a dose-dependent manner (0.1 – 10 μM), resulting in enhanced IFN-γ production, increased frequencies of Granzyme B^+^ CD8^+^ T cells, and reduced AML cell viability (Figure 5E-G for AML blasts and Figure S4D-F for THP-1 cells). These findings demonstrate that pharmacological blockade of LILRB4 by **GL-4512** restores anti-tumor immune function across both solid and hematologic tumor contexts.

Importantly, the ability of **GL-4512** to phenocopy the effects of target-specific anti-LILRB4 antibody blockade across multiple orthogonal functional readouts strongly supports an on-target mechanism of action. Collectively, these results establish **GL-4512** as a functional small-molecule inhibitor capable of reversing SCG2-driven immunosuppressive signaling and restoring anti-tumor immune activity in clinically relevant human co-culture systems.

### In Vitro ADME and In Vivo Pharmacokinetic Profiling Support Once-Daily Dosing of GL-4512 for Efficacy Studies

To evaluate the developability and translational potential of **GL-4512**, we next performed a comprehensive in vitro ADME and safety liability assessment (Table 1). **GL4512** exhibited favorable physicochemical properties, including moderate lipophilicity (LogD_7.4_ = 2.68), good kinetic solubility in aqueous buffer, and sustained solubility under biorelevant FaSSIF conditions. Parallel artificial membrane permeability assay (PAMPA) analysis demonstrated moderate passive membrane permeability (PAMPA *P*_*app*_ = 3.8 × 10^−6^ cm/s), consistent with the compound’s balanced physicochemical profile. **GL-4512** also displayed excellent chemical stability under simulated gastric and intestinal fluid conditions, indicating favorable stability across physiologically relevant gastrointestinal environments (Table 1).

**Table 1.** In vitro PK profile of GL-4512.

| PK parameter | GL-4512 |
| --- | --- |
| LogD<sub>7.4</sub><sup>a</sup> | 2.68 |
| Kinetic Solubility<br>1% DMSO/PBS<br>[$\mu\text{M}$]<sup>b</sup> | 84.7 |
| Solubility in FaSSIF<br>[$\mu\text{M}$] | 71.3 |
| PAMPA $P_{app}$ [cm/s] | $3.8 \times 10^{-6}$ |
| Stability in simulated gastric fluid<br>(pH 1.2, 2h) | >95% |
| Stability in simulated intestinal fluid<br>(pH 6.8, 2h) | >91% |
| $t_{1/2}$ [min] mouse liver microsomes<sup>c</sup> / $CL_{int}$<sup>d</sup> | $96.4 \pm 5.8$ / $18.7 \pm 1.6$ |
| $t_{1/2}$ [min] human liver microsomes<sup>c</sup> / $CL_{int}$<sup>d</sup> | $124.1 \pm 8.1$ / $13.4 \pm 1.2$ |
| $t_{1/2}$ [min] mouse plasma | >180 |
| $t_{1/2}$ [min] human plasma | >180 |
| $t_{1/2}$ [min] PBS | >300 |
| PPB human [%]<sup>c</sup> | $93.1 \pm 0.72$ |
| IC<sub>50</sub> against WI-38<br>[$\mu\text{M}$] | >100 |
| IC<sub>50</sub> against HS 27<br>[$\mu\text{M}$] | >100 |
| IC<sub>50</sub> against hERG<br>[$\mu\text{M}$] | >100 |
| CYP3A4 inhibition<br>(% at 10 $\mu\text{M}$) | 7.1 |
| CYP2D6 inhibition<br>(% at 10 $\mu\text{M}$) | 3.6 |
| CYP2C9 inhibition<br>(% at 10 $\mu\text{M}$) | 5.2 |
| CYP2C19 inhibition<br>(% at 10 $\mu\text{M}$) | 4.4 |

<sup>a</sup>LogD (pH = 7.4) the distribution coefficient at physiological pH 7.4.
<sup>b</sup>Kinetic solubility measured in 1% DMSO/PBS (pH 7.4) by UV–vis spectrophotometry (n=3). <sup>c</sup>Values are mean ± SD (n=3). <sup>d</sup>Intrinsic clearance expressed in $\mu\text{L}/(\text{min} \times \text{mg})$ protein. Values are mean ± SD (n=3).

Metabolic stability studies further demonstrated that **GL4512** possesses a favorable in vitro clearance profile. In mouse and human liver microsomes, **GL-4512** exhibited prolonged half-lives (>90 min) together with relatively low intrinsic clearance values, consistent with moderate metabolic turnover (Table 1). The compound additionally remained highly stable in both mouse and human plasma over the duration of the assay, supporting adequate systemic stability under physiological conditions. Plasma protein binding studies revealed high but acceptable human plasma protein binding, a profile commonly observed among lipophilic immunomodulatory small molecules.

Safety liability profiling demonstrated a clean early developability profile for **GL-4512**. The compound showed no measurable cytotoxicity against normal human fibroblast cell lines (WI-38 and HS-27) at concentrations up to 100 µM. Furthermore, **GL-4512** exhibited negligible inhibition of the hERG potassium channel and minimal inhibition across major cytochrome P450 isoforms, including CYP3A4, CYP2D6, CYP2C9, and CYP2C19, suggesting low potential for cardiotoxicity or CYP-mediated drug–drug interactions. Collectively, these findings indicate that **GL-4512** combines favorable physicochemical properties, metabolic stability, and an acceptable preliminary safety profile suitable for in vivo evaluation.

To determine whether **GL-4512** achieves systemic exposure sufficient to support anti-tumor efficacy studies, in vivo PK studies were performed in C57BL/6 mice following single-dose intravenous (IV) and oral administration. Following IV administration at 2 mg/kg, **GL-4512** exhibited a plasma half-life of 3.8 h, a systemic clearance of 11.2 mL/min/kg, and a volume of distribution of 2.9 L/kg, consistent with moderate clearance and broad tissue distribution. The corresponding plasma exposure (AUC0−∞) was 8.7 µM·h, supporting the favorable in vitro metabolic stability profile observed in microsomal studies.

Following oral administration at 20 mg/kg, **GL-4512** was efficiently absorbed, achieving a plasma C_max_ of 4.9 µM at a T_max_ of 1.2 h. Total systemic exposure following oral dosing reached an AUC0−∞ of 46.3 µM·h, corresponding to an oral bioavailability of approximately 42%. The observed plasma half-life following oral administration was 4.2 h, supporting sustained systemic exposure over the dosing interval. Importantly, plasma concentrations achieved following oral dosing remained substantially above the biochemical and cellular potency ranges observed in TR-FRET competition, SHP1/SHP2 signaling, and p-STAT3 inhibition assays for a significant portion of the dosing interval. These pharmacokinetic characteristics are consistent with effective systemic target coverage following once-daily administration. Taken together, the favorable oral exposure, moderate half-life, acceptable bioavailability, and clean early safety profile of **GL-4512** supported selection of once-daily oral dosing for subsequent efficacy evaluation in the CT26 syngeneic colon carcinoma model.

### GL-4512 demonstrates cross-species LILRB4 engagement and suppresses CT26 tumor progression in vivo

Before evaluating the in vivo anti-tumor activity of **GL4512**, we first assessed whether the compound retained binding activity toward murine LILRB4 to support pharmacological studies in syngeneic mouse tumor models. MST analysis confirmed direct interaction between **GL-4512** and recombinant murine LILRB4, with a dissociation constant K_D_ of 32.5 ± 5.1 nM, indicating that **GL-4512** maintains high-affinity cross-species target engagement suitable for murine efficacy studies (Figure S5). Selectivity profiling using Dianthus/TRIC analysis demonstrated that **GL-4512** produced a strong and selective fluorescence response toward LILRB4 across a panel of 12 related LILR family members and immune checkpoint proteins (Figure S6).

We next investigated the therapeutic activity of **GL-4512** in the CT26 syngeneic colorectal carcinoma model in immunocompetent BALB/c mice. CT26 cells were implanted subcutaneously into female BALB/c mice, and treatment was initiated once tumors reached approximately 75-100 mm^3^. Mice were randomized to receive vehicle or **GL-4512** at 20 mg/kg by oral gavage once daily for 21 consecutive days. The selected dosing regimen was guided by the favorable pharmacokinetic profile of **GL-4512**, which demonstrated sustained systemic exposure above the biochemical and cellular potency range associated with inhibition of LILRB4 signaling.

Daily oral administration of **GL-4512** resulted in significant suppression of CT26 tumor growth compared with vehicle-treated controls, with clear separation of tumor growth curves observed throughout the treatment period (Figure 6A). The anti-tumor activity of **GL-4512** was further reflected by substantial reductions in final tumor weight at study termination (Figure 6B). Importantly, **GL-4512** treatment was well tolerated, with no significant changes in body weight or overt signs of systemic toxicity observed during the study, consistent with the favorable in vitro safety and ADME profile identified during early developability assessment.

**Figure 6.**
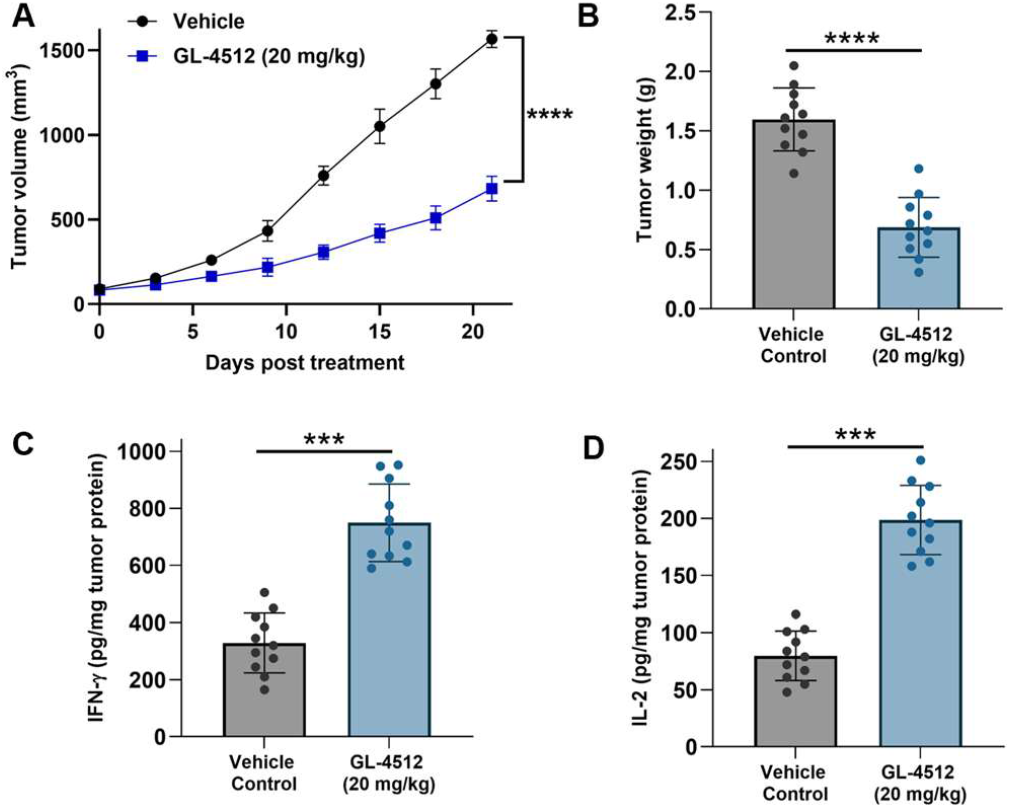
GL-4512 suppresses CT26 tumor progression and enhances anti-tumor immune activity in vivo. **(A)** Tumor growth curves of CT26 syngeneic colorectal tumors in BALB/c mice treated with vehicle or **GL-4512** (20 mg/kg, PO, QD). Treatment was initiated once tumors reached approximately 75–100 mm^3^. **GL-4512** significantly suppressed tumor growth throughout the treatment period. **(B)** Final tumor weights collected at study termination demonstrating significant reduction in tumor burden following **GL-4512** treatment. **(C**,**D)** ELISA analysis of tumor lysates revealed significantly increased intratumoral IFN-γ and IL-2 levels in **GL-4512**-treated tumors relative to vehicle-treated controls, consistent with enhanced anti-tumor immune activation within the tumor microenvironment. Data represent mean ± SEM with individual animals shown. Statistical significance was determined using unpaired two-tailed Student’s t-test. \*\*\**p* < 0.001, \*\*\*\**p* < 0.0001.

To investigate whether the anti-tumor activity of **GL4512** was associated with restoration of anti-tumor immune function in vivo, tumor lysates were analyzed using ELISA-based pharmacodynamic readouts. **GL-4512** treatment significantly increased intratumoral IFN-γ and IL-2 levels relative to vehicle-treated tumors, indicating enhanced immune activation within the tumor microenvironment (Figure 6C,D). These findings are consistent with the ex vivo co-culture studies and support the proposed mechanism whereby pharmacological inhibition of LILRB4 relieves suppressive myeloid signaling and restores productive anti-tumor immune responses in vivo.

## CONCLUSIONS

In this study, we report the discovery and characterization of **GL-4512**, a direct small molecule modulator of the suppressive myeloid immune checkpoint LILRB4 (ILT3). Using a Dianthus/TRIC-based direct target engagement screening workflow, we identified multiple LILRB4-interacting compounds and subsequently validated **GL-4512** through orthogonal biophysical approaches including MST, SPR, and CETSA. Extensive microsecond MD simulations combined with site-directed mutagenesis revealed a previously unrecognized ligandable region within the flexible extracellular domain of LILRB4 and identified W57-associated interactions as important contributors to productive **GL-4512** binding.

Mechanistically, **GL-4512** disrupted the immunosuppressive LILRB4-SCG2 signaling axis and suppressed downstream SHP1/SHP2 and STAT3 signaling pathways associated with myeloid-mediated immune suppression. Importantly, pharmacological inhibition of LILRB4 restored anti-tumor immune activity in clinically relevant colorectal cancer and AML co-culture systems, resulting in enhanced cytokine production, increased cytotoxic T-cell activation, and reduced tumor-cell viability. **GL-4512** additionally demonstrated favorable ADME and pharmacokinetic properties supporting once-daily oral dosing in vivo. In immunocompetent CT26 colorectal tumors, oral administration of **GL-4512** significantly suppressed tumor progression while enhancing intratumoral immune activation. Collectively, these findings establish LILRB4 as a tractable target for direct small molecule immunomodulation and support therapeutic targeting of suppressive myeloid immune checkpoints for cancer using non-biologic modalities.

Beyond the specific context of LILRB4, this work carries broader implications for chemical biology and small molecule drug discovery. The TRIC-based direct engagement screening platform established here is fully applicable to the wider class of non-enzymatic immune checkpoint receptors, including other LILRB family members and additional suppressive myeloid receptors, that have historically resisted small molecule discovery due to the absence of conventional druggable active sites. The identification of a ligandable extracellular pocket through microsecond molecular dynamics simulations and experimental mutagenesis validation further demonstrates that structurally dynamic immune receptors can harbor tractable small molecule binding sites that are not apparent from static structural analysis alone. Collectively, the combination of a generalizable biophysical screening platform, first-in-class direct target engagement, computationally guided binding site identification, patient-derived functional validation across two cancer types, and oral in vivo efficacy reported in a single study establishes a comprehensive framework for future small molecule targeting of suppressive myeloid immune checkpoints. We anticipate that **GL-4512** will serve as a valuable chemical probe for dissecting LILRB4 biology and that the approaches described here will accelerate the discovery of next-generation small molecule cancer immunotherapies targeting the broader landscape of non-enzymatic immune receptors.”

## EXPERIMENTAL

### Dianthus TRIC Assay

Initial screening of the Enamine Hit Locator Library (5,120 compounds) for LILRB4 binding was performed using the temperature-related intensity change (TRIC) assay on a Dianthus NT.23 Pico instrument (NanoTemper Technologies). Recombinant human Histagged ILT3 protein (from SinoBiological, Catalog# 16742-H08H) was fluorescently labeled with RED-tris-NTA 2nd Generation dye (NanoTemper, Cat. #MO-L018) according to the manufacturer’s protocol.

Labeled LILRB4 (final concentration: 30 nM) was incubated with test compounds at a fixed concentration (10 µM) in assay buffer (PBS pH 7.4 supplemented with 0.05% Tween-20 and 1% (v/v) DMSO). Fluorescence was measured in Dianthus 384-well plates (NanoTemper Technologies, Catalog# DI-P001A) with LED power set to 40%. Fluorescence was recorded at 670 nm and 650 nm, and normalized fluorescence (F_norm_) was calculated as the ratio of F670/F650. Binding responses were quantified as Fnorm. Compounds with ΔF_norm_ ≥ 3 units and reproducible responses across replicates (n = 5) were classified as hits.

### Quantitative Binding Affinity Determination Using Monolith

Binding affinities of selected hits were quantified by microscale thermophoresis using a Monolith X instrument (NanoTemper Technologies). His-tagged ILT3 protein was labeled with RED-tris-NTA 2nd Generation dye using the Monolith His-Tag Labeling Kit (Cat. #MO-L018) following the manufacturer’s guidelines. Compound titrations were prepared as serial dilutions (PBS buffer, pH 7.4, 0.05% Tween-20, 1% (v/v) DMSO). Following a 15 min incubation at room temperature in the dark, samples were loaded into Monolith capillaries (Cat. #MO-K022) and analyzed at 25 °C using 40% LED power and medium MST power settings. Normalized fluorescence (F_norm_) values were determined as the ratio of fluorescence intensity after and before IR laser heating. Each compound was evaluated in five technical replicates. Dissociation constants (K_D_) were calculated from three independent experiments using MO.Affinity Analysis software and GraphPad Prism 10, applying standard dose-response fitting models. Data represent mean ± SD (n=5).

### SPR analysis

SPR validation of LILRB4 binding was performed using Biacore™ 8K SPR system (Cytiva, Marlborough, MA, USA). LILRB4-His Protein (10 μg/mL in PBS, pH 7.4) was immobilized on a Series S Sensor Chip CM5 (29104988, Cytiva, Marlborough, MA, USA) using a commercial amine coupling kit (BR100050, Cytiva, Marlborough, MA, USA). Immobilization was performed at a flow rate of 10 μL/min for 420 s, followed by blocking with ethanolamine. The flow cell with solely ethanolamine-block on the same channel served as the reference. Immobilization buffer: PBS-P+ (28995084, Cytiva, Marlborough, MA, USA). Briefly, gradient concentrations of the compound were prepared in the assay buffer, and injected over the sensor chip in a single-cycle kinetics model with a flow rate at 30 μL/min for 120 s per injection. After each injection, a 30 s-regeneration was performed at a flow rate of 30 μL/min using the regeneration buffer. Assays were performed in triplicate, and K_D_ values are reported as mean ± SD (n=3).

### TR-FRET LILRB4–SCG2 Competition Assay

The LILRB4-SCG2 interaction was measured using a homogeneous TR-FRET assay. Recombinant human LILRB4-ECD-Fc and His-tagged SCG2 were incubated with terbium cryptate-labeled anti-His antibody (donor) and XL665-labeled anti-human Fc antibody (acceptor) in medium-binding white 384-well plates (Greiner, #784075). Recombinant human SCG2 carrying C-terminal polyhistidine tag was purchased from SinoBiological (catalog number 13441-H08H), while the ECD of human LILRB4 fused to the Fc region of human IgG1 was obtained from SinoBiological (catalog number 16742-H02H). Terbium cryptate-labeled anti-His monoclonal antibody as the donor fluorophore and an XL665-conjugated anti-human Fc antibody as the acceptor, both sourced from Revvity (catalog numbers 61HISTLF and 61HFCXLF, respectively). Final assay concentrations were 20 nM LILRB4-ECD-Fc, 20 nM SCG2-His, 1 nM donor antibody, and 10 nM acceptor antibody in PBS containing 0.1% DMSO. **GL-4512** was added as a serial dilution and plates were incubated for 2 h at room temperature.

TR-FRET signals were measured using a Tecan Spark plate reader (340 nm excitation; 620 nm donor emission; 665 nm acceptor emission). Data acquisition parameters included 50 flashes per well, a 50 µs delay time, and a 400 µs integration window. A blocking anti-LILRB4 antibody (h128-3, catalog number HV126013) was used as a positive control. Percent inhibition values were normalized to DMSO-treated controls and fitted using nonlinear regression in GraphPad Prism to determine IC_50_ values.

### CETSA

CHO cells stably expressing human ILT3/LILRB4 (from Creative Biogene, Cat# CSC-RO0667) were maintained in Ham’s F-12 medium supplemented with 10% fetal bovine serum and 1% penicillin-streptomycin under standard humidified culture conditions (37 °C, 5% CO2). Cells were seeded in 384-well plates at 12,000 cells/well and incubated overnight at 37 °C in 5% CO_2_. **GL4512** was added at the indicated concentrations and incubated for 60 min. Cells were subsequently subjected to thermal challenge at 51 °C for 3 min (using a calibrated Bio-Rad C1000 Touch thermal cycler), followed by cooling to room temperature. Following thermal challenge, cells were lysed directly in assay plates by addition of lysis buffer supplemented with protease inhibitor cocktail and incubated for 30 min at 4 °C with gentle agitation. Lysates were subsequently clarified by centrifugation at 4000 × g for 10 min to remove aggregated and insoluble protein species generated during thermal denaturation.

The soluble fraction of LILRB4 remaining after thermal challenge was quantified using an AlphaLISA-based immunodetection format. Briefly, clarified lysates were incubated with ILT3-specific biotinylated detection antibody and AlphaLISA acceptor beads according to the manufacturer’s instructions, followed by addition of streptavidin-coated donor beads under reduced-light conditions. After incubation at room temperature, AlphaLISA signal was measured using Tecan Spark plate reader. CETSA stabilization signals were normalized to vehicle-treated controls, and dose-response curves were generated using nonlinear regression analysis. Data represent mean ± SD, n = 5.

### Computational studies

#### LILRB4 Modeling

The globular domain of LILRB4 structure (residues P17-S206) was obtained from the AF3 model corresponding to the UniProt sequence. The available crystal structure (PDB ID: 3P2T) was not used due to an engineered disulfide bond introduced to stabilize the structure and aid crystallisation.^48^ **GL-4512** coordinates were constructed from its SMILES representation using UCSF Chimera.^49^

#### Molecular Dynamics System Preparation

The ILILRB4 AF3 model and two molecules (Mol1 and Mol2) of **GL-4512** were parameterized using the CHARMM36^50^ force field for the protein and the CGenFF^51^ force field for the ligand. The system was solvated in a periodic box filled with TIP3P water molecules,^52^ with a minimum padding of 15 Å from the protein surface. System preparation was performed using in-house scripts that combined Python-based HTMD^53^ workflows with Tool Command Language (Tcl) scripts. Protonation states were assigned using pdb2pqr and PROPKA3^54^, assuming a physiological pH of 7.4. Solvation was performed using the VMD Solvate plugin (version 1.5), and charge neutrality was achieved by adding Na^+^ and Cl^−^ ions to reach a final ionic strength of 0.15 M using the VMD Autoionize plugin (version 1.3).

#### MD Simulations

All simulations were performed using the ACEMD3 engine.^55^ The systems were first equilibrated under isothermal–isobaric (NPT) conditions at 310 K and 1 atm. Temperature was controlled using a Langevin thermostat^56^ with a damping coefficient of 1 ps^−1^, while pressure was maintained using a Monte Carlo barostat.^57^ An integration time step of 2 fs was used during equilibration. Initial steric clashes were relieved by 500 steps of conjugate-gradient energy minimization. This was followed by an equilibration phase, during which positional restraints of 1 kcal mol^−1^ Å^−2^ were applied to protein atoms and gradually released. Production simulations were then carried out in the canonical ensemble (NVT). Four independent replicas of 5 μs each were generated, resulting in a total aggregated simulation time of 20 μs. These simulations employed a 4 fs integration time step, with temperature maintained at 310 K using a Langevin thermostat with a damping coefficient of 0.1 ps^−1^. Bond lengths involving hydrogen atoms were constrained using the M-SHAKE algorithm.^58^ Non-bonded interactions were computed using a cutoff distance of 9 Å, with a switching function applied from 7.5 Å. Long-range electrostatics were treated using the particle mesh Ewald (PME) method^59^ with a grid spacing of 1.0 Å.

#### MD Trajectories Analysis

Trajectory analyses were carried out using VMD^60^ and complementary tools. Root mean square deviation (RMSD) and root mean square fluctuation (RMSF) were computed using VMD. MMGBSA analysis was performed using the MMPBSA.py script^61^ from the Amber-Tools20 suite,^62^ considering 1 μs of simulations for Site 1 (Replica 1) and Site 2 (Replica 4). VMD VolMap plugin 1.1 was employed to compute the volumetric maps of the two molecules of compound 2, with a grid resolution of 1 Å, on the full aggregated trajectory corresponding to 20 μs of simulation time.

#### THP-1-Derived Macrophage-Like Cell Signaling Assays

THP-1 cells (from ATCC) were seeded at 1 × 10^6^ cells/well in 6-well plates and differentiated into macrophage-like cells by treatment with phorbol 12-myristate 13-acetate (PMA, 100 nM) for 48 h, followed by a 24 h resting period in PMA-free complete RPMI-1640 medium supplemented with 10% fetal bovine serum and 1% penicillin– streptomycin. Differentiated cells were stimulated with recombinant human SCG2 (100 ng/mL) in the presence of increasing concentrations of **GL-4512** for 1 h at 37 °C. Cells were subsequently washed with ice-cold PBS and lysed using RIPA buffer supplemented with protease and phosphatase inhibitor cocktails. Lysates were clarified by centrifugation at 14,000 × g for 15 min at 4 °C prior to downstream phospho-signaling analysis.

Phosphorylated SHP1, phosphorylated SHP2, and phosphorylated STAT3 were quantified using commercial phosphospecific ELISA kits (abcam, catalog# ab279924, ab314344, and ab279941, respectively) according to the manufacturers’ protocols. Lysates were normalized for total protein content using a BCA assay before loading. Data were normalized to SCG2-treated controls and expressed as percent of SCG2-induced signaling. IC_50_ values were calculated using nonlinear regression. Data represent mean ± SD, n = 5.

#### Human Co-Culture Assays

For colorectal cancer co-culture assays, HCT116 or HT-29 cells (from ATCC) were seeded in flat-bottom 96-well plates at 1 × 10^4^ cells/well and allowed to adhere overnight in complete RPMI-1640 medium supplemented with 10% fetal bovine serum and 1% penicillin–streptomycin. PBMCs isolated from colorectal cancer patients (from STEMCELL Technologies) by Ficoll density-gradient centrifugation were added at an effector-to-target ratio of 10:1 in the presence or absence of recombinant human SCG2 (100 ng/mL) and increasing concentrations of **GL-4512** (0.1–10 μM). A blocking anti-LILRB4 antibody (h128-3, catalog number HV126013) was included as a target-specific positive control. Co-cultures were incubated for 72 h at 37 °C in 5% CO_2_. Supernatants were collected for IFN-γ and IL-2 quantification by ELISA (abcam catalog# ab46025 and #ab270883, respectively), and tumor-cell viability was assessed using CellTiter-Glo (from Promega) according to the manufacturer’s protocol.

For AML co-culture assays, THP-1 cells (from ATCC) or primary AML blasts (from STEMCELL Technologies) were co-cultured with PBMCs from AML patients (from STEMCELL Technologies) or enriched CD3^+^ T cells under analogous conditions. Supernatants were collected for IFN-γ and IL-2 ELISA analysis (abcam catalog# ab46025 and #ab100706, respectively). Cytotoxic T-cell activation was assessed by measuring Granzyme B^+^ CD8^+^ T cells using flow cytometry. Tumor-cell viability was quantified using CellTiter-Glo (from Promega). Data were normalized to media or SCG2-treated controls as indicated in the figure legends. Data represent mean ± SD, n = 5.

#### Site-directed mutagenesis

Alanine substitutions (W57A and Y192A) were introduced into the human ILT3 extracellular domain using the QuikChange Lightning Site-Directed Mutagenesis Kit (Agilent Technologies) according to the manufacturer’s protocol. Briefly, mutagenic primers were designed to incorporate the desired codon substitutions and PCR amplification was performed using a high-fidelity polymerase. Parental methylated DNA was digested with DpnI, and the resulting plasmids were transformed into *E. coli* DH5α cells. All mutations were confirmed by Sanger sequencing. Recombinant wild-type and mutant ILT3 proteins were expressed and purified as described above. Protein integrity and purity (>90%) were verified by SDS-PAGE and analytical size-exclusion chromatography. Binding affinities of mutant proteins were evaluated by MST under identical conditions to wild-type controls, and Kd values were obtained from nonlinear regression fits as described above.

#### PK studies

In vitro PK profiling was performed as we previously reported.^40,41^ In vivo PK studies were conducted in male C57BL/6 mice (8-10 weeks old; n = 5 per time point). The tested compound was administered by intravenous injection (2 mg/kg) or oral gavage (20 mg/kg) in a formulation consisting of 5% DMSO, 40% PEG400, and 55% sterile saline (v/v/v), prepared fresh prior to dosing. Blood samples were collected at 0.25, 0.5, 1, 2, 4, 8, and 24 h postdose via tail vein sampling into EDTA-coated tubes. Plasma was isolated by centrifugation at 3,000 × g for 10 min at 4°C.

#### CT26 Syngeneic Colorectal Tumor Model

All animal studies were performed in accordance with protocols approved by the Institutional Animal Care and Use Committee (IACUC). Female BALB/c mice (6-8 weeks old) were obtained from Jackson Laboratory and acclimated for at least 1 week prior to experimentation under standard pathogen-free housing conditions with ad libitum access to food and water.

CT26 murine colorectal carcinoma cells were maintained in RPMI-1640 medium supplemented with 10% fetal bovine serum and 1% penicillin–streptomycin. Cells were harvested during logarithmic growth phase, washed twice with sterile PBS, and resuspended in cold PBS. Each mouse received a subcutaneous injection of 5 × 10^5^ CT26 cells into the right flank in a final injection volume of 100 µL. Tumor growth was monitored three times weekly using digital calipers. Once tumors reached approximately 75-100 mm^3^, mice were randomized into treatment groups (n = 11 mice/group) to ensure comparable mean starting tumor volumes across cohorts. **GL-4512** was formulated fresh daily in vehicle consisting of 5% DMSO, 40% PEG400, and 55% sterile saline (v/v/v). Mice received vehicle control or **GL-4512** at 20 mg/kg by oral gavage once daily for 21 consecutive days. The selected dosing regimen was guided by in vivo PK studies demonstrating sustained plasma exposure above the biochemical and cellular potency range associated with LILRB4 signaling inhibition.

Body weight and clinical condition were monitored throughout the study to assess tolerability. Humane endpoints included tumor ulceration, impaired mobility, or tumor burden exceeding institutional limits. At study termination, mice were euthanized by CO_2_ inhalation followed by cervical dislocation. Tumors were excised, weighed, photographed, and snap-frozen for downstream pharmacodynamic analyses.

Tumor lysates were prepared using ice-cold lysis buffer supplemented with protease inhibitors and clarified by centrifugation. Total protein concentrations were quantified using a BCA assay prior to normalization. Intratumoral IFN-γ and IL-2 levels were quantified using commercial ELISA kits (abcam catalog# 100689 and #ab46096, respectively) according to the manufacturers’ protocols. Cytokine levels were normalized to total tumor protein content and expressed as pg/mg tumor protein.

#### Statistical Analysis

Data are presented as mean ± SD or mean ± SEM as indicated in the figure legends. Statistical analyses were performed using GraphPad Prism. For two-group comparisons, unpaired two-tailed Student’s t-tests were used. For multiple-group comparisons, one-way ANOVA followed by Tukey’s post hoc test was applied. Tumor growth curves were analyzed using repeated-measures ANOVA where appropriate. *P* values <0.05 were considered statistically significant.

## ASSOCIATED CONTENT

### Supporting Information

The Supporting Information is available free of charge on the ACS Publications website.

Chemical structures of primary hit compounds, validation of binding site by site-directed mutagenesis, MST dose response curves, co-culture assays of PBMCs and cancer cells (PDF)

## AUTHOR INFORMATION

### Author Contributions

The manuscript was written through contributions of all authors. All authors have given approval to the final version of the manuscript.

## ABBREVIATIONS

ADME: absorption, distribution, metabolism, and excretion
AF3: AlphaFold3
AML: acute myeloid leukemia
BLI: biolayer interferometry
CETSA: cellular thermal shift assay
ECD: extracellular domain
ELISA: enzyme-linked immunosorbent assay
IL-2: interleukin-2
ILT3: immunoglobulin-like transcript 3
ITIM: immunoreceptor tyrosine-based inhibitory motif
KD: equilibrium dissociation constant
LILRB4: leukocyte immunoglobulin-like receptor B4
MD: molecular dynamics
MDSCs: myeloid-derived suppressor cells
MST: microscale thermophoresis
PBS: phosphate-buffered saline
PBMCs: peripheral blood mononuclear cells
PK: pharmacokinetics
PPB: plasma protein binding
RMSD: root mean square deviation
RMSF: root mean square fluctuation
SCG2: secretogranin II
SD: standard deviation
SEM: standard error of the mean
SHP1: Src homology region 2 domain-containing phosphatase-1
SHP2: Src homology region 2 domain-containing phosphatase-2
SPR: surface plasmon resonance
STAT3: signal transducer and activator of transcription 3
TAMs: tumor-associated macrophages
THP-1: human monocytic leukemia cell line
TIME: tumor immune microenvironment
TR-FRET: time-resolved fluorescence resonance energy transfer
TRIC: temperature-related intensity change
WI-38: human lung fibroblast cell line.

## Notes

The authors declare no competing financial interests.

## Table of Contents artwork

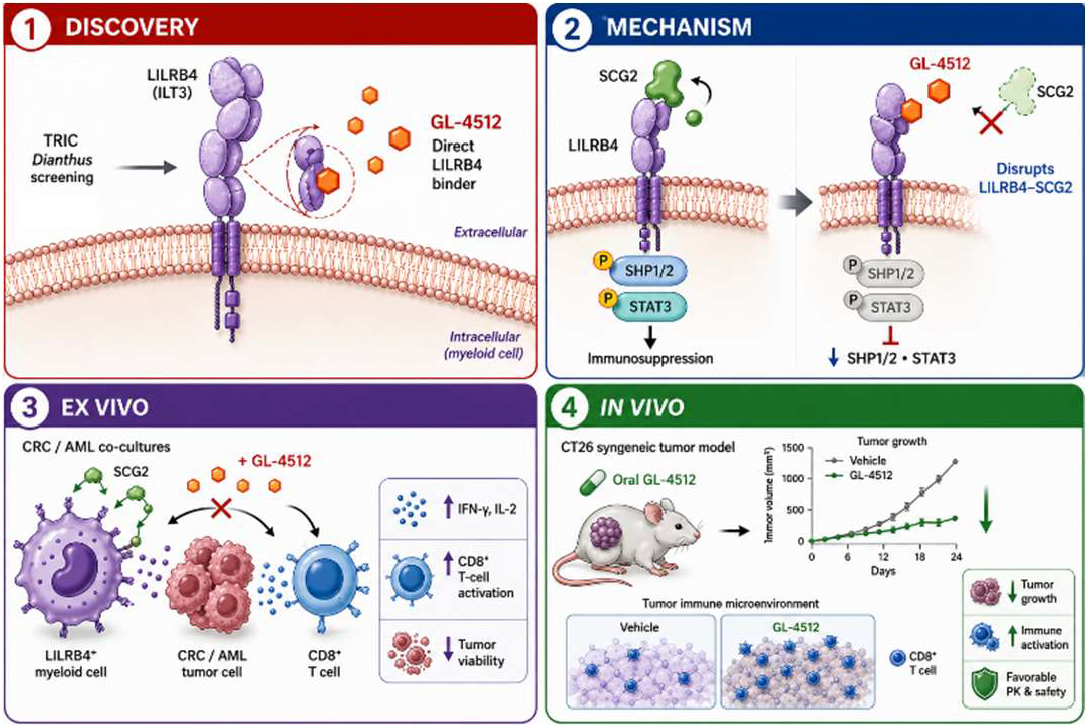

