## Supporting Information for "Direct Small Molecule Modulation of LILRB4 (ILT3) Restores Anti-Tumor Immunity In Vivo and in Patient-Derived Cells"

*Electronic Supplementary Information*

| **Contents** | |  |
| --- | --- | --- |
| Chemical structures of the validated hit compounds identified by Dianthus screening | | S2 |
| MST analysis of **GL-2047** binding to recombinant human LILRB4 | | S3 |
| MST analysis of **GL-3901** binding to recombinant human LILRB4  Site-directed mutagenesis supports Site 2 as the dominant **GL-4512** binding pocket on LILRB4 | | S3  S4 |
| **GL-4512** restores anti-tumor immune activity across colorectal cancer and AML co-culture systems  **GL-4512** directly binds recombinant murine LILRB4 as determined by MST  Selectivity profiling of **GL-4512** across immune checkpoint and LILR family proteins | | S5  S6  S7 |

**Table S1**. Chemical structures of the validated hit compounds identified by Dianthus screening.

| **Compound name** | **Chemical structure** | **LILRB4 (K_D_)** |
| --- | --- | --- |
| **GL-4512** | 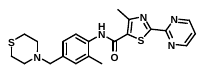 | 18.1 ± 2.4 nM |
| **GL-2047** | 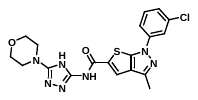 | 503 ± 19.2 nM |
| **GL-3901** | 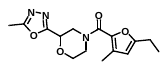 | 3.18 ± 0.65 μM |
| **GL-0884** | 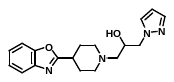 | 8.32 ± 1.04 μM |
| **GL-1228** | 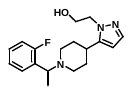 | 12.43 ± 3.29 μM |
| **GL-3015** | 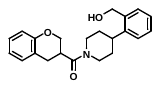 | 19.29 ± 4.58 μM |

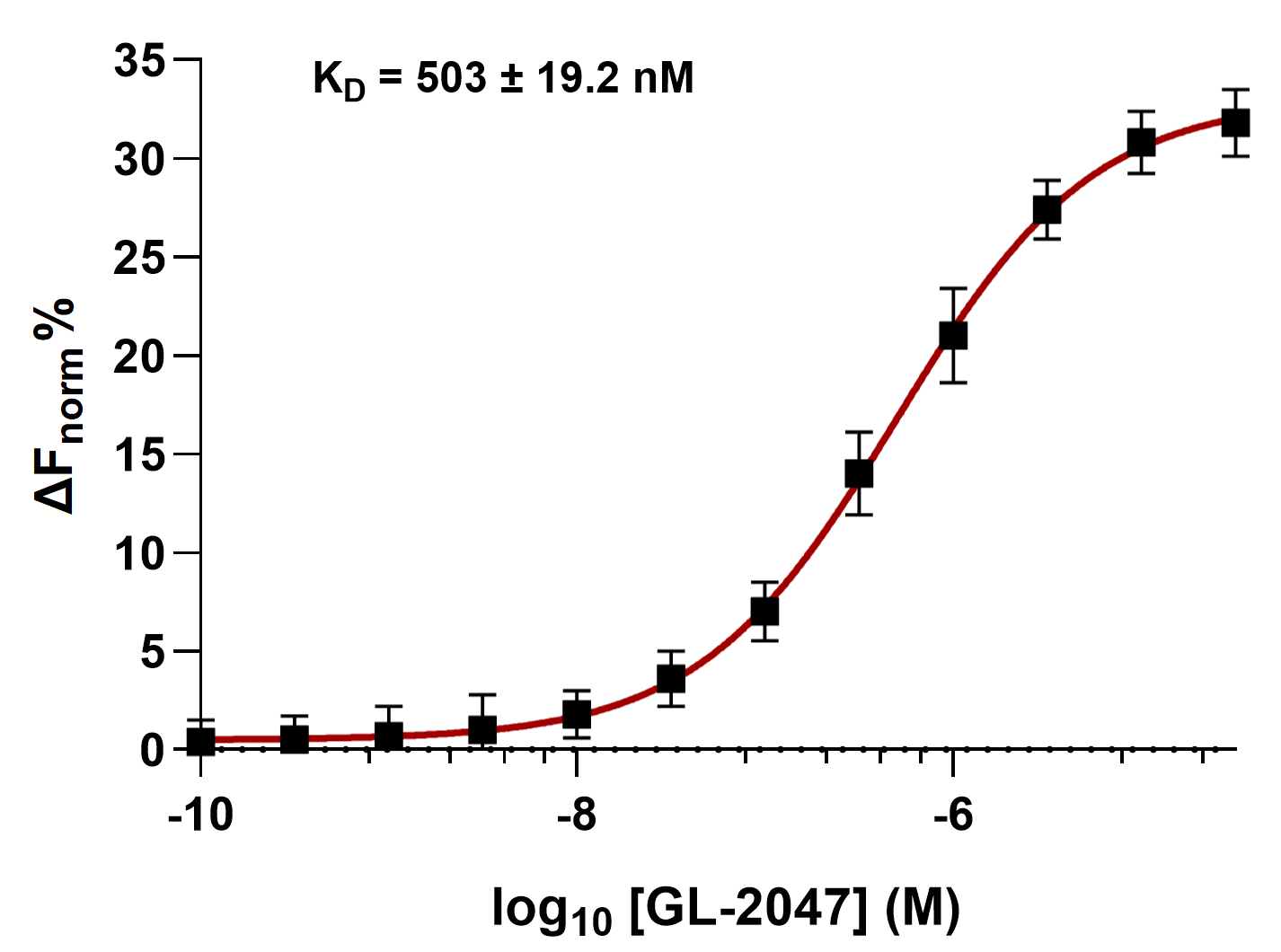

**Figure S1. MST analysis of GL-2047 binding to recombinant human LILRB4.** Microscale thermophoresis (MST) analysis demonstrating concentration-dependent binding of **GL-2047** to recombinant human LILRB4 extracellular domain protein. Changes in normalized fluorescence (ΔF_norm_) were used to quantify target engagement. Data represent mean ± SD, n = 5.

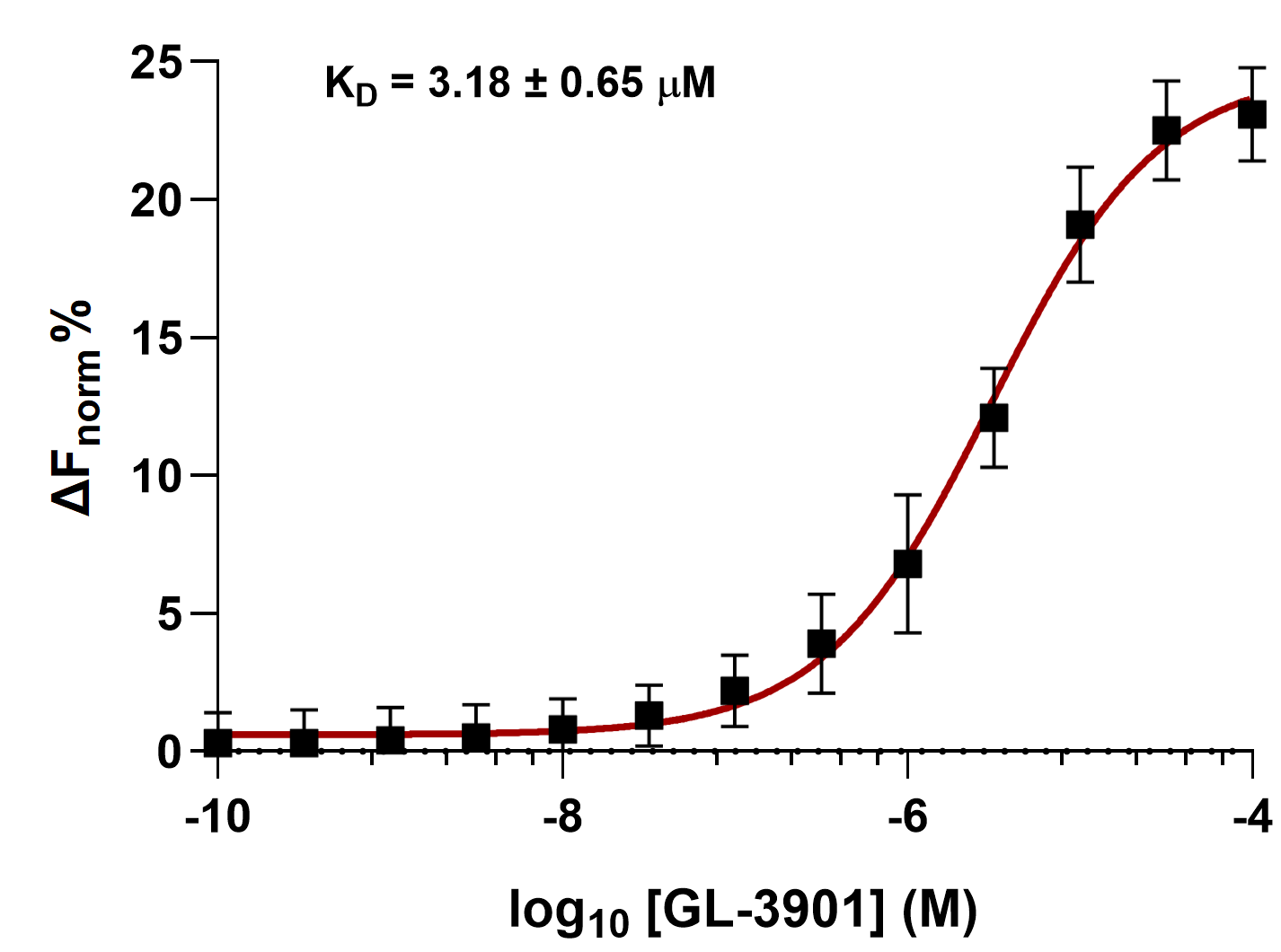

**Figure S2. MST analysis of GL-3901 binding to recombinant human LILRB4.** Microscale thermophoresis (MST) analysis demonstrating concentration-dependent binding of **GL-2047** to recombinant human LILRB4 extracellular domain protein. Changes in normalized fluorescence (ΔF_norm_) were used to quantify target engagement. Data represent mean ± SD, n = 5.

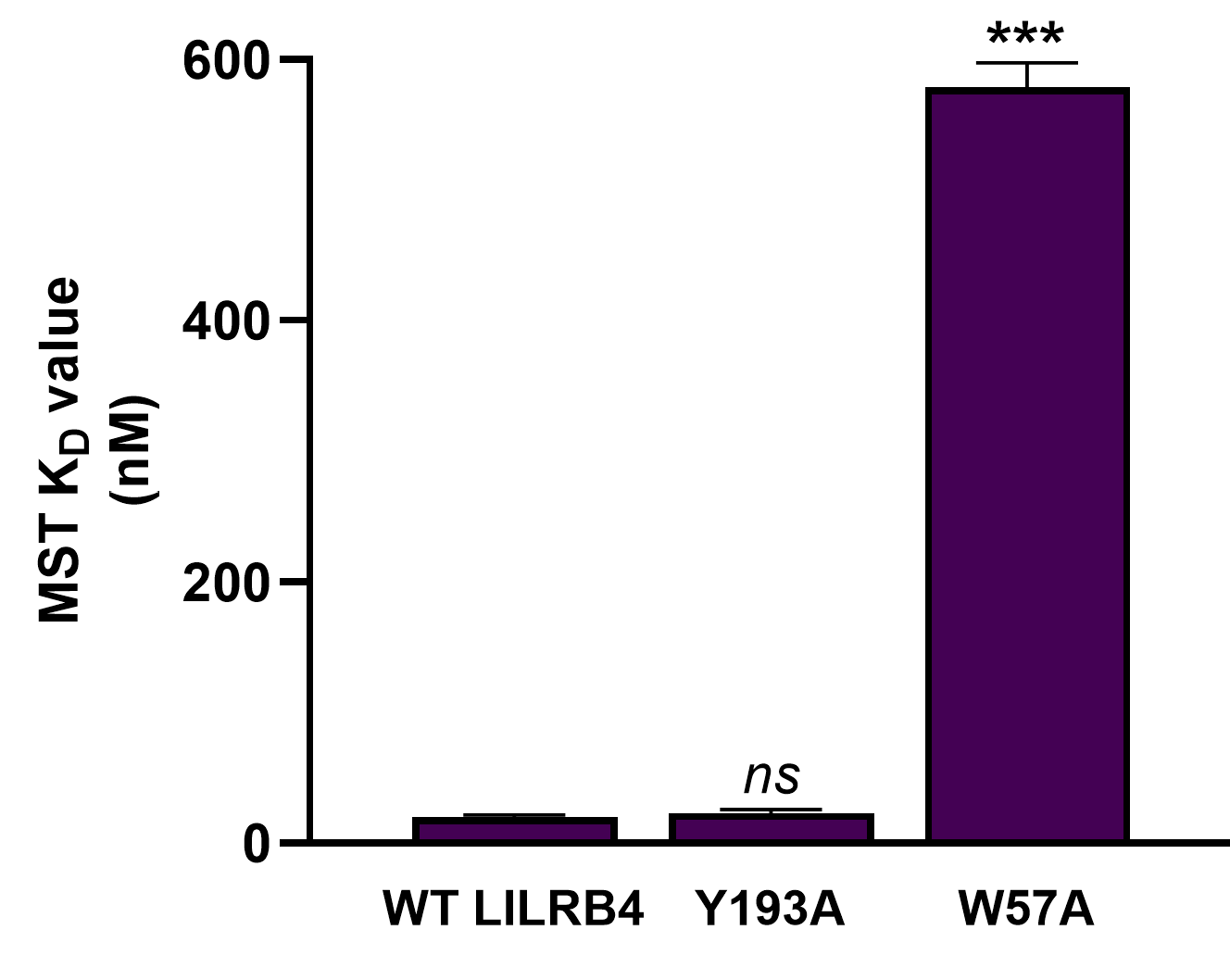

**Figure S3. Site-directed mutagenesis supports Site 2 as the dominant GL-4512 binding pocket on LILRB4.** MST analysis was performed to evaluate the contribution of representative residues from the two computationally predicted **GL-4512** binding sites on LILRB4. Mutation of Y193, a residue associated with the transiently sampled Site 1 interaction region, produced no significant effect on **GL-4512** binding affinity relative to wild-type (WT) LILRB4. In contrast, mutation of W57, a key aromatic residue within the energetically favorable Site 2 pocket, resulted in a marked reduction in binding affinity, producing an approximately 32-fold increase in the apparent equilibrium dissociation constant (KD). These findings support Site 2 as the principal productive binding pocket mediating high-affinity interaction of **GL-4512** with LILRB4 and experimentally validate the molecular dynamics-predicted binding model. Data represent mean ± SD, n = 5. Statistical significance was determined using one-way ANOVA followed by post hoc multiple comparison testing; *ns*, not significant; ****p* < 0.001.

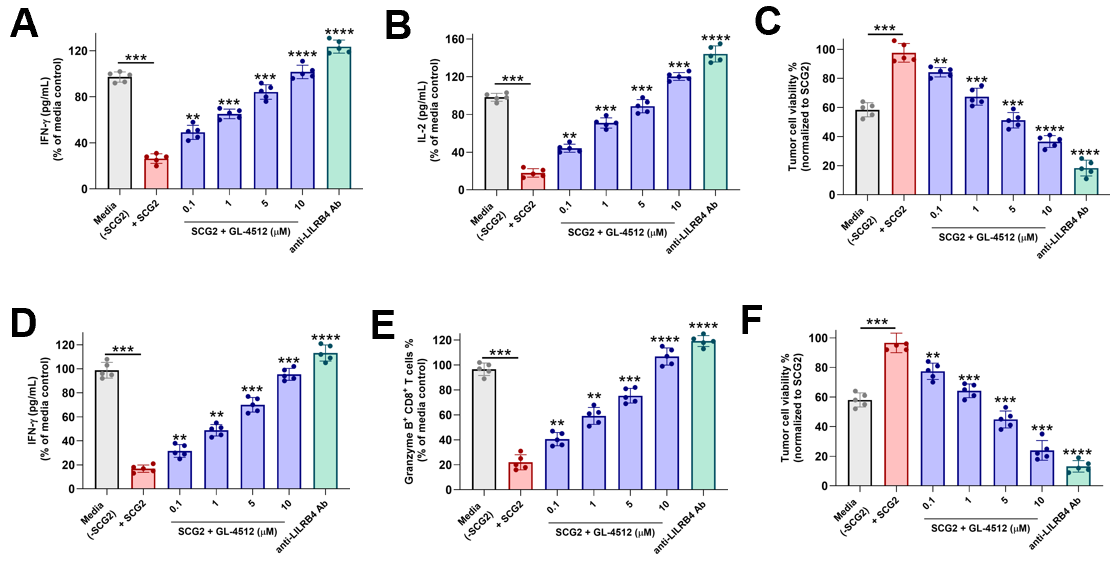

**Figure S4. GL-4512 restores anti-tumor immune activity across colorectal cancer and AML co-culture systems. (A,B)** In colorectal HT-29 cancer co-cultures, SCG2 stimulation suppressed anti-tumor immune activity, resulting in reduced IFN-γ and IL-2 secretion. Treatment with **GL-4512** restored cytokine production in a concentration-dependent manner. **(C) GL-4512** significantly reduced colorectal HT-29 cancer cell viability in SCG2-treated co-cultures, consistent with restoration of immune-mediated tumor killing. **(D)** In THP-1 AML co-cultures, **GL-4512** restored IFN-γ secretion suppressed by SCG2-mediated LILRB4 signaling. **(E) GL-4512** increased the frequency of Granzyme B^+^ CD8^+^ T cells in a dose-dependent manner, indicating recovery of cytotoxic T-cell function. **(F) GL-4512** significantly reduced THP-1 AML viability in SCG2-treated co-cultures. Data represent mean ± SD from independent donors performed in technical replicates. Statistical significance was determined using one-way ANOVA with Tukey’s multiple comparisons test. ***p* < 0.05, ****p* < 0.001, *****p* < 0.0001 relative to media +SCG2.

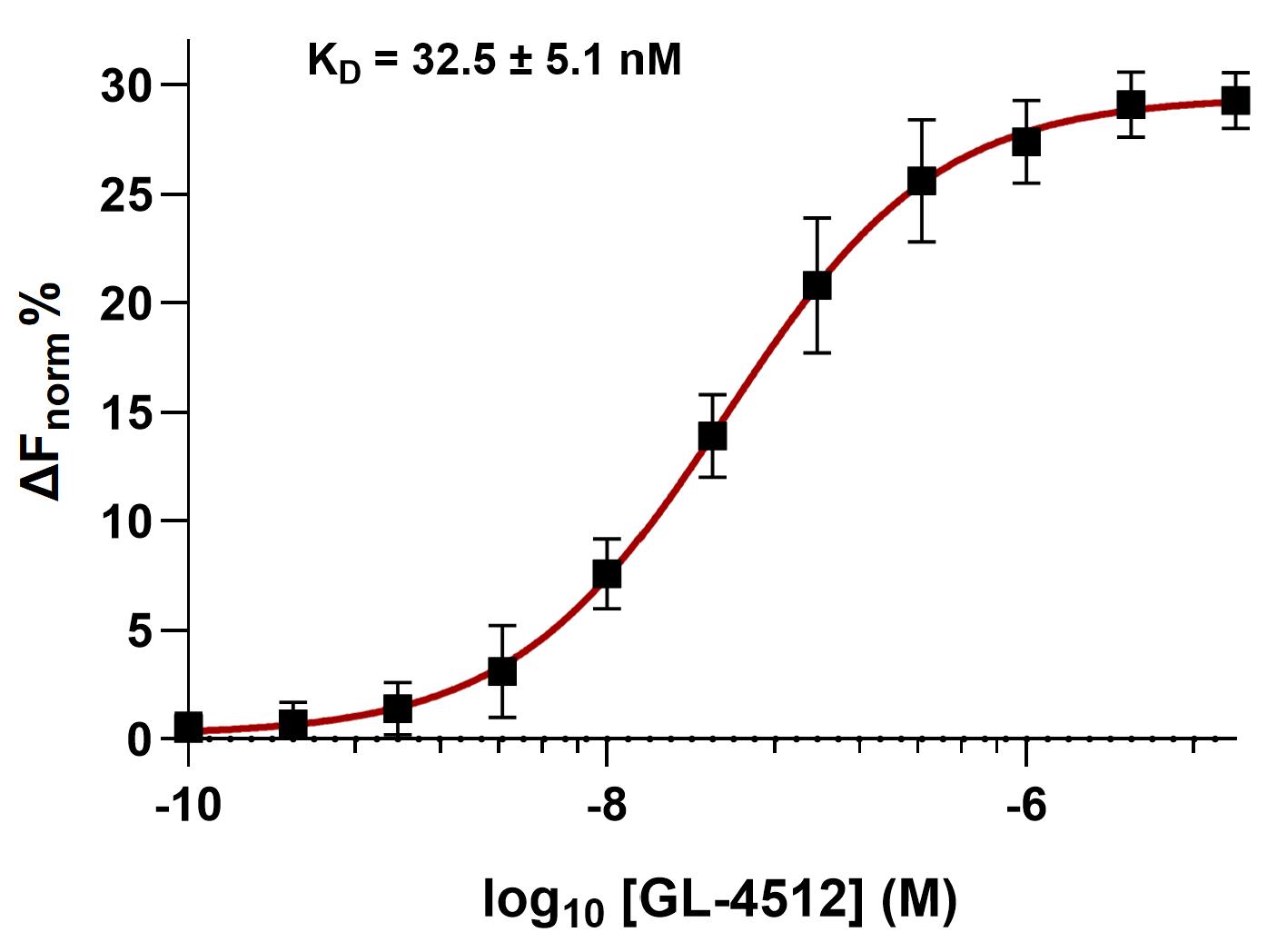

**Figure S5. GL-4512 directly binds recombinant murine LILRB4 as determined by MST.** Binding of **GL-4512** to recombinant murine LILRB4 was evaluated using MST under solution-phase conditions. **GL-4512** exhibited concentration-dependent interaction with murine LILRB4 with a calculated dissociation constant (KD) of 33.7 nM, demonstrating high-affinity cross-species target engagement suitable for evaluation in syngeneic mouse tumor models. Data represent mean ± SD from three independent measurements.

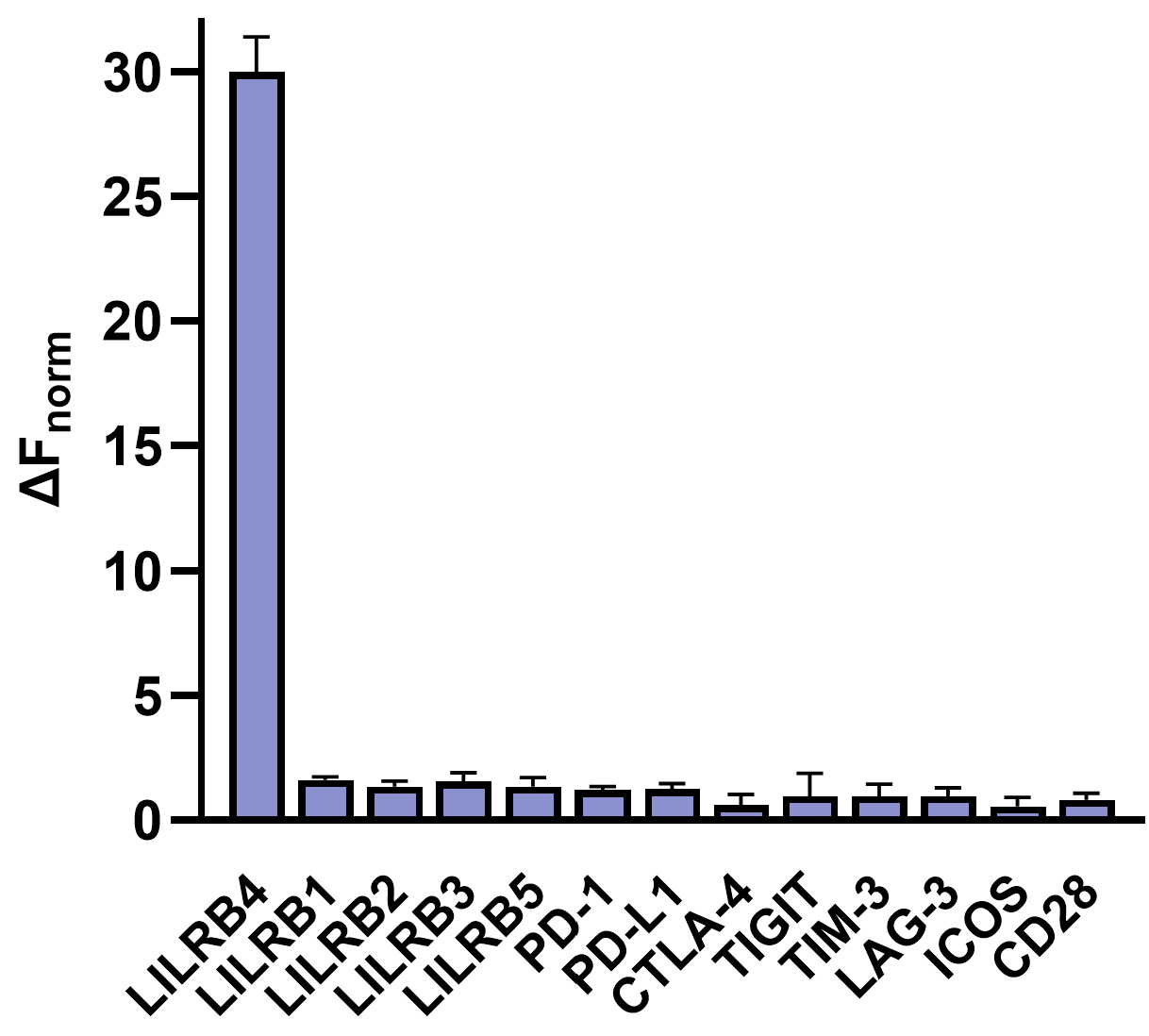

**Figure S6. Selectivity profiling of GL-4512 across immune checkpoint and LILR family proteins.**
Dianthus/TRIC analysis showing normalized fluorescence changes (ΔF_norm_) following incubation of **GL-4512** with a panel of immune checkpoint proteins and related LILR family members. **GL-4512** produced a strong and selective ΔF_norm_ response toward LILRB4 (ILT3), while minimal responses were observed across related LILRB homologs and canonical immune checkpoints including PD-1, PD-L1, CTLA-4, TIGIT, TIM-3, LAG-3, ICOS, and CD28. Data represent mean ± SD (n = 5).
